# Anther cones increase pollen release in buzz-pollinated *Solanum* flowers

**DOI:** 10.1101/2021.10.02.462843

**Authors:** Mario Vallejo-Marín, Carlos Eduardo Pereira Nunes, Avery Leigh Russell

**Affiliations:** Biological and Environmental Sciences. University of Stirling, Stirling, FK9 4LA, Scotland, United Kingdom; Department of Biology, Missouri State University, Springfield, Missouri 65897. U.S.A.

**Author notes:** **Author contributions:** MVM, CEPN, and ALR conceived the idea. MVM designed and performed the experiments, wrote the software, and analysed the data. CEPN collected and analysed data. MVM, CEPN, and ALR wrote the manuscript and contributed to reviewing and editing it. MVM acquired the funding. **Data accessibility:** The vibration and pollen data will be made publicly available in DataSTORRE (https://datastorre.stir.ac.uk) upon publication.

**Keywords:** Anther cone, bees, buzz pollination, convergent evolution, pollen release, pollination

## Abstract

The widespread evolution of tube-like anthers releasing pollen from apical pores is associated with buzz pollination, in which bees vibrate flowers to remove pollen. The mechanical connection among anthers in buzz-pollinated species varies from loosely held conformations, to anthers tightly held together with trichomes or bio-adhesives forming a functionally joined conical structure (anther cone). Joined anther cones in buzz-pollinated species have evolved independently across plant families and via different genetic mechanisms, yet their functional significance remains mostly untested. We used experimental manipulations to compare vibrational and functional (pollen release) consequences of joined anther cones in three buzz-pollinated species of *Solanum* (Solanaceae). We applied bee-like vibrations to focal anthers in flowers with (“joined”) and without (“free”) experimentally created joined anther cones, and characterised vibrations transmitted to other anthers and the amount of pollen released. We found that joined anther architectures cause non-focal anthers to vibrate at higher amplitudes than free architectures. Moreover, in the two species with naturally loosely held anthers, anther fusion increases pollen release, while in the species with a free but naturally compact architecture it does not. We discuss hypotheses for the adaptive significance of the convergent evolution of joined anther cones.

## Introduction

Flowers are extremely morphologically diverse, and establishing how this morphological diversity affects function has long been a focus of research (Darwin 1877; Vogel 1996). Buzz-pollinated plants capture the close relationship between floral form and function. In these species, modifications of floral structures result in morphologies that require the visits of bees that produce vibrations to remove pollen grains (Macior 1968; Thorp and Estes 1975; Buchmann et al. 1977). The floral vibrations produced by the bee cause the anthers to shake, transmitting energy to the pollen grains inside the anthers and causing them to be propelled outwards through the apical pores (Buchmann and Hurley 1978). In buzz-pollinated plants, floral structures, usually the anthers, but sometimes the corolla, have evolved a tubular form that retains the pollen grains inside after anthesis (Buchmann 1983; Vallejo-Marín 2019). A taxonomically widespread floral form of buzz-pollinated plants that has evolved convergently across multiple plant families is the *Solanum*-like or “solanoid” flower, named by Fægri (1986) after the canonical flower form of *Solanum* (Solanaceae) (Endress 1996a; De Luca and Vallejo-Marin 2013; Russell et al. 2016). In *Solanum-*like flowers, the anthers are often arranged in the centre of the flower forming a structure that resembles a cone (Faegri 1986). The degree to which the anthers in the solanoid flower are physically connected to one another varies. In one extreme, the enlarged stamens might be held loosely towards the centre of the flower, with each individual stamen capable of relatively independent movement from the other stamens (“free” stamen architecture). Other species may have connivent anthers that are closely pressed together yet non-joined. In the other extreme, anthers can be physically attached to each other (postgenitally connate; Endress 1996a), forming a single conical structure (Glover et al. 2004). This type of joined anther cone (“joined” architecture) has evolved multiple independent times in different plant groups including species in the families Anthericaceae, Luzuriageaceae, Pittosporaceae, Tecophilaceae and Solanaceae (Endress 1996a; Glover et al. 2004). Despite the repeated evolution of conical anther architecture across different species, to date, no studies have tried to empirically evaluate its functional significance.

During buzz pollination, an individual bee might only vibrate one or few anthers. However, the vibrations generated by the bee’s thorax and applied to this focal anther(s), propagate through the flower (Brito et al. 2020; Nevard et al. 2021), and can cause pollen release even in distal anthers that are not in direct contact with the bee’s body (Arroyo-Correa et al. 2019). These oscillations in the focal anther(s) can cause other stamens to vibrate via two transmission pathways: **(1) Filament pathway**. Anthers are attached to the corolla or to the base of the flower via a filament that, in buzz pollinated plants is usually short and relatively stiff. The oscillation of the focal anther can thus cause shaking in the whole flower via the filament attachment, which in turn causes vibrations in other, non-focal anthers. **(2) Anther-anther pathway**. In species with floral architectures in which stamens are held closely together, for example forming a cone (joined anthers) (Glover et al. 2004), vibrations can be transmitted by direct anther-anther contact from the vibrating focal anther to adjacent anthers even when these distal anthers are not touching the bee’s body. Recent work across buzz-pollinated flowers with different morphologies in three different plant families (Solanaceae, Primulaceae and Gentianaceae), suggests that stamen architecture, defined as the stamen’s relative sizes, degree of fusion and their spatial and functional connections (Endress 1996b), affects the transmission of vibrations (Nevard et al. 2021). Therefore, variation in stamen architecture could be associated with different types of vibrations experienced by distal, non-focal anthers during buzz pollination with potential consequences for pollen release and pollen placement on the pollinator’s body (Glover et al. 2004; Nevard et al. 2021).

Here we use an experimental approach to compare the vibrational and functional (pollen release) consequences of joined anther cones in buzz-pollinated species in the genus *Solanum* (Solanaceae). Specifically, we address the following two questions: **(1)** How do the vibrations experienced in stamens differ between floral configurations with free vs. joined anthers? We hypothesise that when vibrations are applied to a focal anther (proximate anther), the vibration amplitude experienced by distal anthers (those anthers not directly being vibrated) is higher in floral configurations with joined anthers than in floral configurations with free anthers. Our hypothesis assumes that species with loose anthers mainly transmit vibrations to distal anthers via the filament pathway, while anther fusion enables vibration transmission via both the filament and anther-anther pathways. **(2)** How does anther fusion into a cone affects pollen release upon vibrations? We hypothesise that the higher vibration amplitude of anthers in joined architectures result in higher pollen release compared to free anther configurations. Our hypothesis is based on the fact that higher vibration amplitudes (e.g., higher velocity or acceleration amplitude) have been shown to be theoretically (Buchmann and Hurley 1978; Hansen et al. 2021) and empirically associated with higher rates of pollen release (De Luca et al. 2013; Rosi-Denadai et al. 2020; Kemp and Vallejo-Marin 2021). If joined anther architectures are associated with higher vibration magnitudes across more anthers (both focal and distal anthers), then we would expect pollen release to be proportionally higher as well.

## Material and Methods

### Study system

*Solanum* is the largest buzz-pollinated genus of flowering plants with approximately 1,400 species (Knapp 2002; Särkinen et al. 2013). Within the genus *Solanum* L. and its close relatives, joined anther cones have evolved independently at least four times: in tomatoes (*S. lycopersicum* L., sect. *Lycopersicon*) and its wild relatives, in *S. dulcamara* L. (sect. *Dulcamara*), in *S. luridifuscescens* Bitter (sect. *Cyphomandropsis*) and related taxa (Glover et al. 2004; Falcāo et al. 2016), and in some species of *Lycianthes* such as *L. synanthera* and *L. anomala* (Dean et al. 2020). Strikingly, the joined anther cone in clades of *Solanum* is formed via different attachment mechanisms. In *S. lycopersicum* and *S. luridifuscescens*, the anther cone is formed via interlocking epidermal cells, joining adjacent anthers, while in *S. dulcamara* smooth anthers are held together by adhesive secretions (Glover et al. 2004; Falcāo et al. 2016). In some cases, the pores of individual anthers in species with joined cones can secondarily evolve increasingly longitudinally dehiscent slits as in *S. lycopersicum* and related taxa. In these species, the slits open to the interior of the joined cone which functions as a single poricidal unit (Endress 1996a).

We studied three *Solanum* species from the subgenus Leptostemonum that differ in stamen architecture, specifically, the extent to which the anthers are loosely or closely held together: *Solanum sisymbriifolium* Lam. and *S. elaeagnifolium* Cav. have free, relatively loose, stamen architectures, while *S. pyracanthos* Lam. has stamens that, although not joined, are held closely together forming a cone-like structure. In *S. sisymbriifolium* and *S. elaeagnifolium*, the petals become reflexed soon after anthesis, and the anthers, which are loosely connivent at anthesis, become increasingly spread out (Knapp 2014a; Vorontsova and Knapp 2014). In *S. pyracanthos*, the petals are slightly reflexed, and the stamens are free but held closely together forming a conical structure that persists throughout anthesis. All three species are nectarless, andromonoecious (producing both hermaphroditic and staminate flowers in the same individual) (D’Arcy 1992; Knapp 2014a; Vorontsova and Knapp 2014), and present flowers to pollinators more or less horizontally, with the anthers’ long axis pointing parallel to the ground (Figure 1). We used seeds of two accessions of *S. sisymbriifolium*, either from seeds collected in the field and outcrossed in the glasshouse or sourced from a commercial provider (Chiltern Seeds, Wallingford, UK) (Table 1). For *S. elaeagnifolium* we used two accessions of this species collected in arid regions in Mexico where they formed locally abundant populations along roadsides and train tracks (Table 1). A summary of the distribution, floral characteristics, and floral visitors of these species as well as the accessions used in this study is presented in Table 1.

**Table 1.**
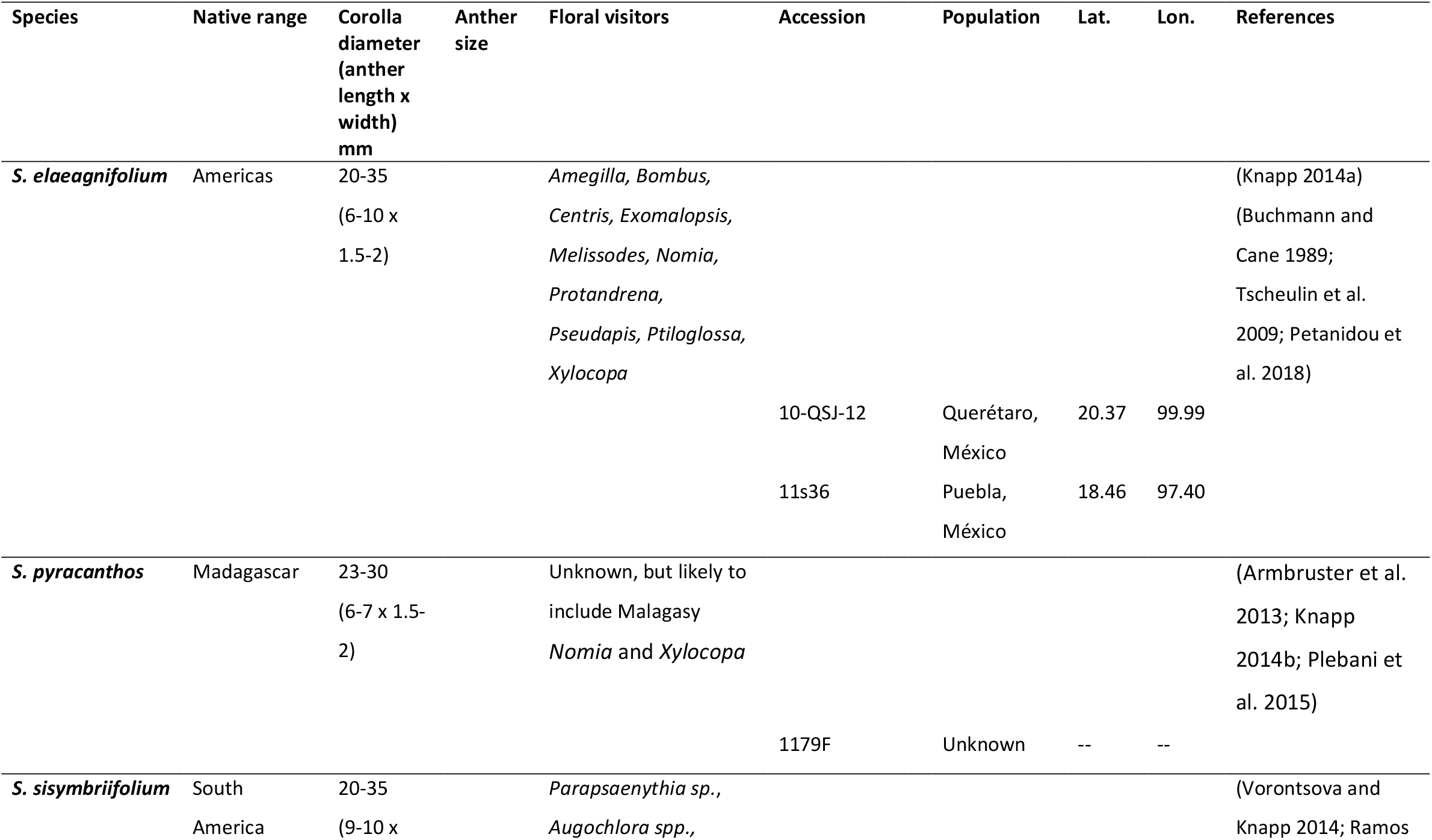
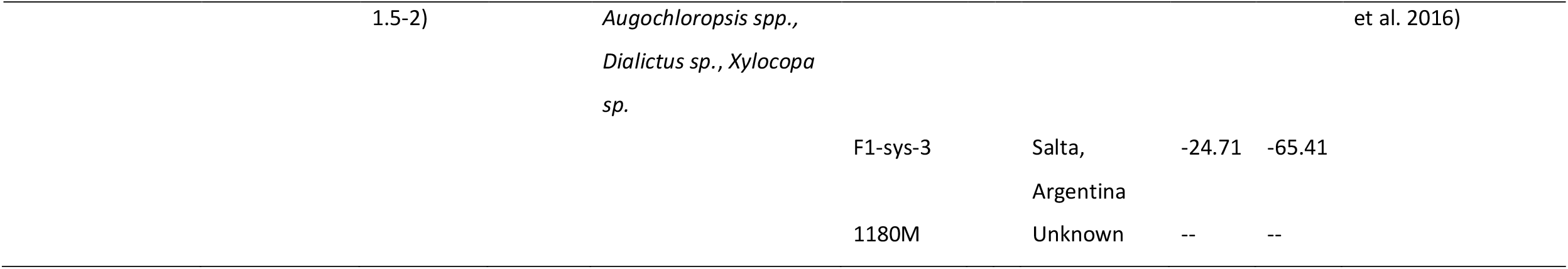
Species distribution, flower and anther characteristics, floral visitors and accessions of the three *Solanum* species used in the present study. Accessions 1179F and 1180M were sourced commercially (see Methods) and their exact origin is unknown.

**Figure 1.**
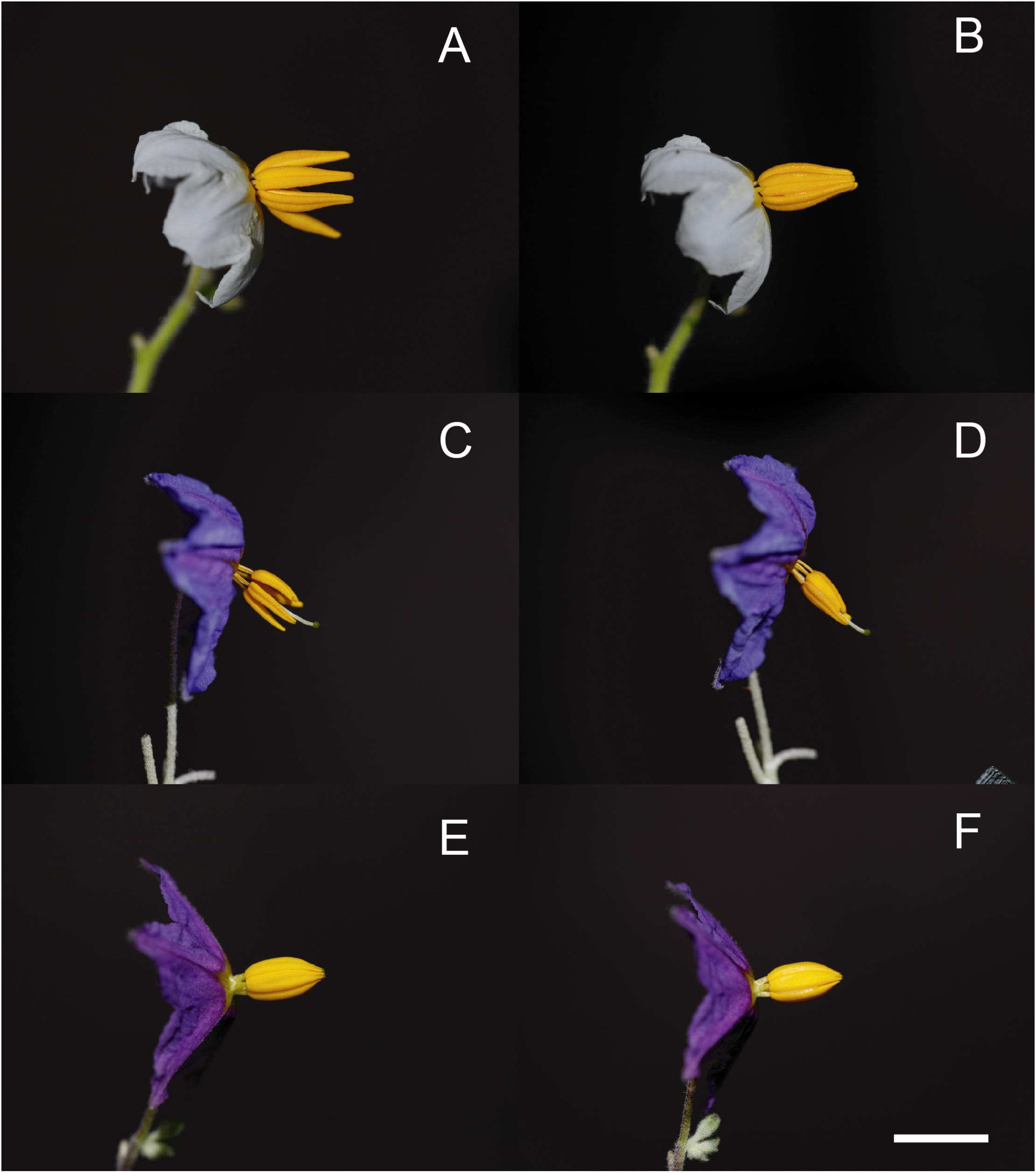
Flower profiles of the three studied species of *Solanum* before (left-hand side panels) and after (right-hand side panels) applying the experimental treatment joining the anthers in a joined cone. The left hand-side panels show the flowers in their natural form and orientation. Notice the natural variation in the level of contact between the individual anthers across the three species, with *S. sisymbriifolium* and *S. elaeagnifolium* having relatively free anthers, while the anthers of *S. pyracanthos* are naturally arranged in a connivent cone. A, B: *S. sisymbriifolium;* C, D: *S. elaeagnifolium*; E, F: *S. pyracanthos*. Scale bar = 1cm.

### Plant growth

Seeds were germinated following a 24h treatment with 1000ppm aqueous solution of gibberellic acid (GA3, Asklepios-seeds, Bad Liebenzell, Germany) at room temperature. Approximately 10 days after germination, seedlings were transplanted to 1.5L plastic pots with an 80:20 compost mix of John Innes No. 2 and medium grade perlite (LBS Horticulture, Colne, UK), and placed in a Snijders Microclimate cabinet in the Controlled Environment Facilities at the University of Stirling with the following growth conditions: 16h-light/8h-dark cycles, at 27°C/25°C, with constant 50% RH. A few additional plants of *S. pyracanthos* and *S. elaeagnifolium* were transplanted to 10L pots and placed in the glasshouse with 16h of supplemental light using compact fluorescent lamps and provided with heating to keep temperature above 16C at night. Plants were fertilised as needed with Tomorite Concentrated Tomato Food (Levington, Surrey, UK).

### Signal generation

To mimic a vibration produced by bees during buzz pollination, we synthesised a pure tone at 300Hz of two seconds in duration with a 50ms fade-in and fade-out feature using *seewave* (Sueur et al. 2008) and saved it as a Mono WAV file at a sampling rate of 44.1kHz. The frequency and duration of the synthetic buzz is within the range observed in buzz pollinating bees (De Luca and Vallejo-Marin 2013; De Luca et al. 2019). The 50ms fade-in and fade-out features in the synthesised signal were introduced to avoid amplitude spikes in the playback equipment caused by a rapid voltage change.

### Vibration system

We built a vibration system to generate and apply vibrations to experimental flowers (Figure 2). A permanent magnetic shaker (LDV210, Brüel & Kjær, Nærum, Denmark) attached to a linear power amplifier (LDS-LPA100, Brüel & Kjær) that received the signal played back in the computer was used to generate the vibration. An M4, 10cm stainless steel screw (“wand”) (ACCU, Huddersfield, UK) was attached to the magnetic shaker. A miniature IPC force sensor (209C11, PCB Piezotronics) was placed at the end of the wand. The system was attached to the flower using an entomological pin “00” of 0.30mm in diameter (Austerlitz, Entomoravia, Czech Republic) cut 10 mm from the tip and superglued to a head screw in the force sensor (Figure 2).

**Figure 2.**
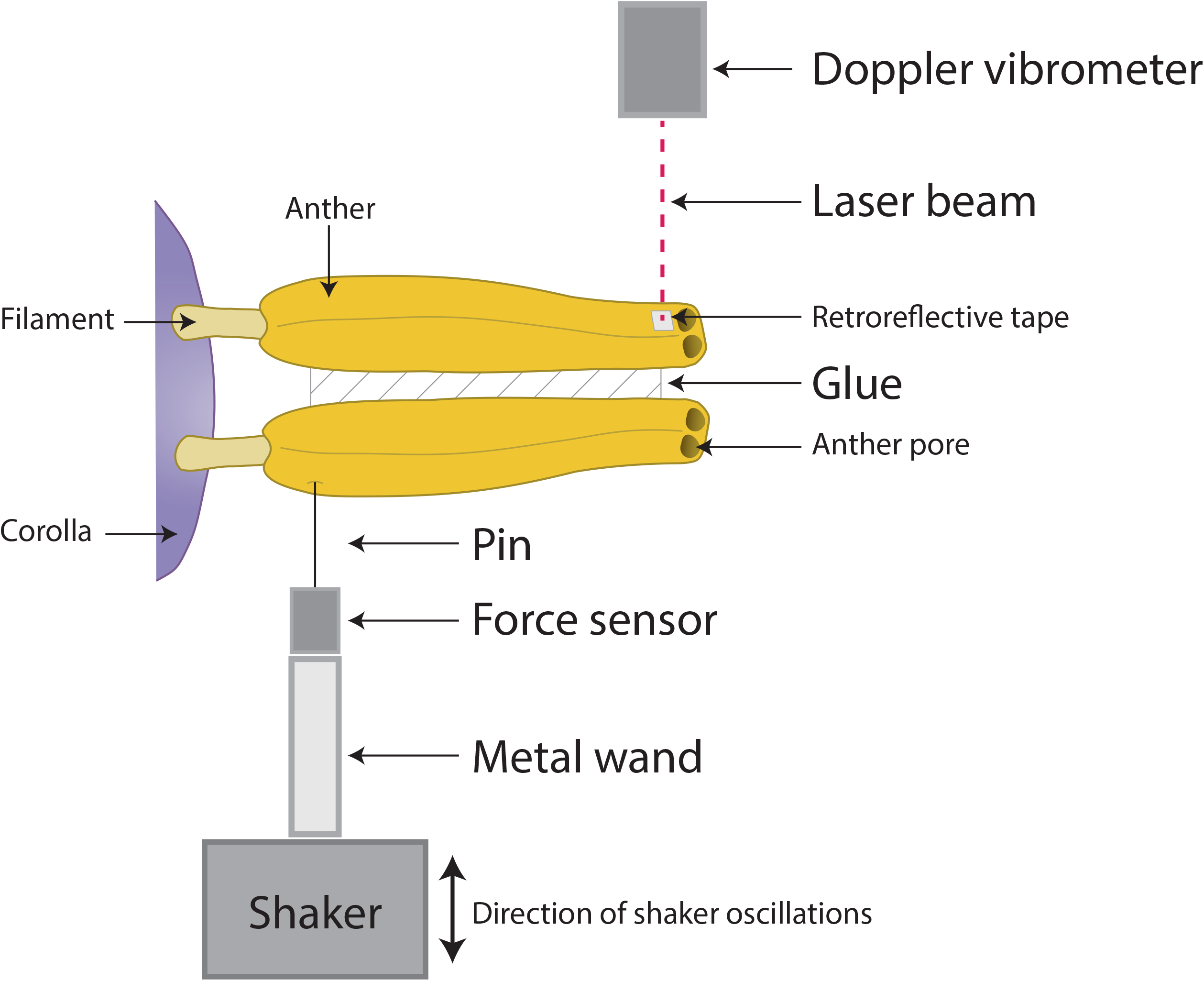
Diagram of the experimental set up (for details see Methods). Diagram not to scale.

### Vibration playback

The synthesised signal was played back in Audacity ver. 3.0 (Audacity 2021) using a laptop computer (HP Elitebook) with the volume set to 70% and output to the linear amplifier. We adjusted the gain in the linear power amplifier by hand to generate approximately 80mm/s peak velocity (∼56mm/s, Root Mean Square (RMS) velocity) measured in a small piece of retroreflective tape at the base of the pin. The tape was kept in place after calibration. The value of peak velocity was chosen to be within the range of values observed in buzz-pollinating bumblebees (De Luca et al. 2013), and used previously in pollen release experiments (Kemp and Vallejo-Marin 2021). Calibration measurements were taken and recorded each day at the start and end of the experimental run.

### Experimental treatment

Flowers were collected in the morning from plants growing in the cabinets or, more rarely, the glasshouses and immediately transferred to a temperature- and humidity-controlled laboratory (19C, 40% RH) for data acquisition. The age of the flower was recorded as days since anthesis (day 1 = flowered opened that day). Flowers were removed from the inflorescence by cutting at the base of the pedicel and placing them in wet floral foam (Oasis Ideal Floral Foam Maxlife, Smithers-Oasis UK Ltd, Washington, UK) on a plastic container. Flowers were measured within 1-3 hours of being removed from plant.

Once in the lab, each flower was randomly assigned to one of two treatments: (1) **Free configuration** This represents the natural arrangement of the flowers. A small amount of PVA glue was applied as a sham treatment to the outside of the anther. (2) **Joined configuration** The anthers were glued together using a small amount of PVA applied along the lateral edge of each anther without blocking the pores. The treatment was applied with the help of a dissecting scope (6.7x magnification). Every flower in the experiment experienced both treatments assigned in a random order. When the free configuration treatment was applied to a flower that had been previously glued for the joined configuration treatment, the anthers were freed using fine-tip forceps and carefully separating the anthers from each other while leaving the glue on the anther (in some cases a small amount of glue fell off in the process of freeing the anthers).

To increase the reflectivity of the anther surface for Doppler vibrometry, we placed a small square of retroreflective tape (approximately 1-4mm^2^) onto a single anther in the adaxial side of the flower (anther 3 or 4 in Figure 3) depending on which one presented a surface that was perpendicular to the laser beam and parallel to the axis of the vibrations produced by the shaker (Figure 2). The tape was placed as close to the tip of the anther as possible, without blocking the pores. The tape was attached to anthers 3 or 4 (at the top of the flower). The shaker pin was inserted at the base of anther 1 (the lowest most anther, see Figure 3). Sometimes, we used the dissecting microscope to draw a small dot made with a black marker to help placing the pin in the exact desired location. The pin made a microscopic wound in the anther. The pin was carefully pushed into the anther without going all the way through.

**Figure 3.**
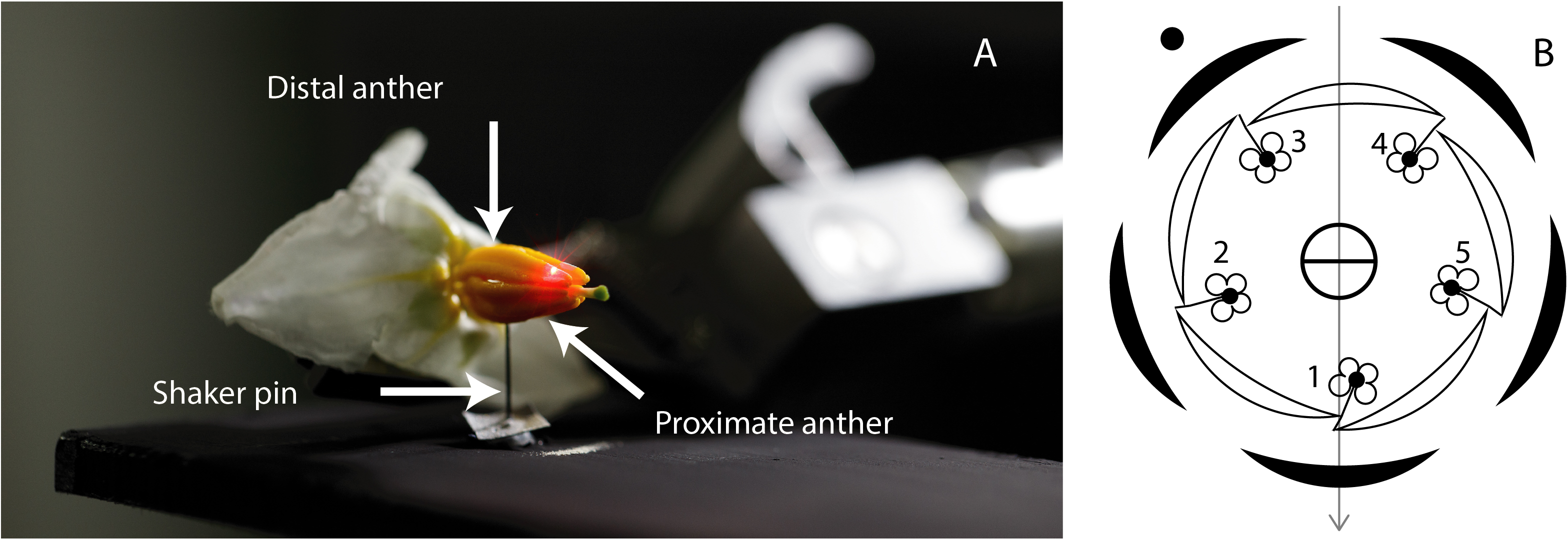
(**A**) Close up of the experimental set up showing the attachment of the shaker system via an entomological pin to the proximate anther (anther 1 in B) and the laser beam of the Doppler vibrometer targeting one of the distal anthers (either anther 3 or 4 in B). **(B)** Floral diagram of *Solanum* showing the labelled anthers (1-5). Floral diagram modified from (Robyns 1931; Knapp 2002).

During the experiment, the flower was held by the pedicel using a stainless-steel micro-V clamp (VK250/M; Thorlabs Ltd., Ely, UK) on a 100mm post (TR100/M), held in a 40mm post holder (PH40/M), perpendicular to the ground. The post holder was screwed in an adjustable-angle mounting plate (AP180/M; Thorlabs) placed on a linear stage (M-UMR8.25; Newport Spectra-Physics Ltd, Didcot, UK) with a standard resolution micrometer (BM17.25; Newport) (see Jankauski 2020). The system was attached to a vertical translation stage (VAP10/M; Thorlabs) fixed to a 250×300×12.7mm aluminium breadboard (MB2530/M). To reduce external vibration noise, the breadboard was placed on four sorbothane isolators (25.4mm × 27mm in diameter; AV4/M) and rested on an antivibration table.

### Vibration recording

To measure the velocity of the vibrations applied and measured, we used a Doppler laser vibrometer (PDV-100, Polytec Ltd, Coventry, UK) set to 500mm/s maximum velocity and a Low Pass Filter at 22kHz. The force applied by the shaker was simultaneously measured using the miniature force sensor. The signals of both the laser vibrometer and the force sensor were simultaneously acquired using a two-channel NI9250 Sound and Vibration module (NI Corporation (UK) Ltd, Newbury, UK) and a USB powered data acquisition module (cDAQ-9171, NI). The acquisition was done using custom-written software in LabView NXG 5.1 (NI). Samples were acquired at a rate of 10,240 samples per second. Data was saved in TDMS format and subsequently converted to text files using a custom program in LabView.

### Signal processing

For each recorded vibration, we estimated the dominant frequency (Hz) and the RMS amplitude for both the velocity (mm/s) measured in the distal anther, and the force (mN) measured in the proximate anther. The text file containing the velocity and force measurements for each vibration was processed in R ver. 4.0.5 (R Core Development Team 2021) using the package *seewave* (Sueur et al. 2008). In brief, we first removed the offset of the signal and used the *timer* function (threshold = 3, dmin = 1, window size = 30, no overlap) on the force channel to identify the segment of the recording to be analysed (approximately 2 seconds or 20,480 samples per channel). For each channel, we then applied a high pass filter at 20Hz (Hanning window, window length = 520 samples) to remove background noise. The filtered signal was then used to obtain a power spectrum using the function *spec* (PSD = TRUE) with a Hanning window of 1,024 samples and a frequency range of 0-2,000Hz. The RMS amplitude was calculated on the same filtered signal.

### Pollen collection

The pollen ejected through floral vibrations was collected on a plate made of black polyethylene measuring 13 cm × 5cm × 4mm (4083829; RS Components Ltd, Corby, UK) (see Ito et al. 2020). The plate had a hole drilled at 2cm from one of the short edges at the midline to allow the shaker wand to go through. The surface of the plate was painted with black acrylic paint (Black 3.0, Culture Hustle, Dorset, UK) (2 layers) to increase the contrast of pollen grains against the background. The polyethylene slide was positioned immediately below the flower with the aid of a micromanipulator (M330 with M3 tilting base and 2.5kg weight; World Precision Instruments, Hitchin, UK). After the vibration was applied, the ejected, light-coloured, pollen fell on the slide and provided a good contrast against the black background. The pollen was collected from the slide using a 2mm^3^ cube of fuchsine-glycerol jelly kept at room temperature. The jelly with the collected pollen was then placed in a 1.5mL microcentrifuge tube and stored at 4°C until pollen counting.

### Pollen counting

To estimate the number of pollen grains released after experimental buzz, we used a particle counter (Multisizer 4e Coulter Counter, Indianapolis, USA). Each pollen sample (contained in a 2mm^3^ piece of fuchsin jelly in a vial) was suspended in 1 ml distilled water by heating it at 80°C for at least 30 min then shaking the vial vigorously with the help of an electric shaker until the vial content looked homogeneous. Prior to counting with the particle counter, the vial contents were added to 19 ml 0.9% NaCl solution, for a total volume of approximately 20 mL. For each sample, the amount of pollen was counted in four 1 ml subsamples. Only particles within the size range of 15-30μm were included in downstream analysis. Although the size of viable pollen grains within the studied species is much less variable and averaged around 24-25μm (Supplementary Figure 1), using a broader particle size range allowed us to also include inviable pollen grains, which were considerable smaller. In total, the particle analysis counted and measured 2,291,835 particles in the 15-30μm range (Supplementary Figure 1). We ran blank samples, containing only 0.9% NaCl solution at the beginning of every daily session and regularly between samples to ensure the equipment calibration accuracy. Finally, we summed the pollen counts for the four subsamples and multiplied by five to obtain each pollen release estimate.

### Statistical analysis

To analyse the effect of anther treatment (free vs. joined) on vibration RMS amplitude we used generalized mixed effects models using the package *lme4* (Bates et al. 2014) in *R* ver. *4*.*0*.*5* (R Core Development Team 2021). We fitted separate models for RMS velocity of the distal anther and for RMS force of the proximate anther. For each response variable, we started by fitting a full model with anther fusion treatment (free vs. joined), plant species, the interaction between anther treatment and plant species, sequential buzz order (first or second), and flower age (in days) as fixed effects. Plant accession, individual, and flower identity were used as random effects to yield the following model:

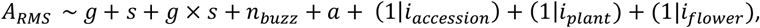

where *A*_*RMS*_ is vibration RMS amplitude (either velocity or force), *g* is anther fusion treatment, *s* is plant species, *n*_*buzz*_ is sequential buzz order, *a* is flower age, and *i*_*accession*_, *i*_*plant*_, and *i*_*flower*_ are indices corresponding to accession, individual and flower identity, respectively. For distal anther velocity, the full model yielded a singular fit and thus we reduced model complexity by removing the plant accession random effect (*i*_*accession*_). This simplified model yielded qualitatively identical results to the full model (results not shown) while avoiding the fit singularity. For proximate anther velocity, the full model also yielded a singular fit, and thus we sequentially removed the random effects of plant accession and plant identity (*i*_*accession*_, *i*_*plant*_). The simpler model also yielded qualitatively identical results (coefficients and statistical significance) to the full model (results not shown). No overdispersion of the residuals of the final models was detected with the statistical package *DHARMa* (Hartig 2021).

To test for the effect of anther configuration on pollen release, we also fitted a generalized mixed effects model using the number of pollen grains removed per buzz as the response variable. The response variable was square root transformed to improve the distribution of the residuals. The fitted model was:

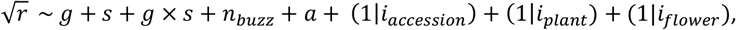

where *r* is number of pollen grains removed, and other variables are as previously defined. No singular fit was detected in this case and the full model was used in the final analysis. No overdispersion of the residuals was detected with the statistical package *DHARMa* (Hartig 2021). Statistical significance of fixed effects (*P-*values) for all final models were obtained using Type III analysis of variance and Satterthwaite’s estimation of degrees of freedom implemented in the package *lmerTest* (Kuznetsova et al. 2017). Given that we detected species *x* treatment interactions (see Results), we calculated the estimated marginal means with treatment nested within species using the fitted mixed-effects model in the *R* package *emmeans* (Lenth 2021). Statistical significance of the linear contrast (free – joined) was obtained using a Kenward-Roger approximation of degrees of freedom.

## Results

We found a statistically significant effect of anther fusion treatment, which depended on plant species, on both the velocity of the distal anther and on the force measured in the proximate anther (treatment *x* species interaction; Table 2, Figure 4). For distal anther velocity, we found that anther fusion increased the RMS velocity achieved by distal anthers compared to the free anther treatment (Figure 4). The effect of treatment on RMS velocity was significant for all within species comparisons (estimated marginal means contrasts, *P* < 0.0001) but the marginal means contrast was 73-83% higher for *S. elaeagnifolium* (110.0 ± 8.93 mm/s) and *S. sisymbriifolium* (116.4 ± 6.88mm/s) than for *S. pyracanthos* (63.3 ± 8.58 mm/s) (Figure 4). For both *S. elaeagnifolium* and *S. sisymbriifolium*, the RMS velocity at the tip of the distal anther in the free treatment was lower than the input RMS velocity measured in the shaker, while the anther velocity in the joined treatment was higher than the input velocity for all species (Figure 5). For the force measured in the proximate anther, we also found a species-dependent effect of treatment (Table 2). All species experienced a higher force in the proximate anther in the joined treatment (estimated marginal means contrasts, *P* < 0.005), although the magnitude of this difference varied across species (marginal means difference = 2.43 ± 0.827 mN, 2.54 ± 0.637 mN, 6.93 ± 0.795 mN, for *S. elaeagnifolium, S. sisymbriifolium* and *S. pyracanthos*, respectively) (Figure 4). In *S. elaeagnifolium* and *S. pyracanthos*, the force measured in the proximate anther was generally higher than the input force measured in the shaker before loading the flower, but in *S. sisymbriifolium* RMS force in the proximate anther varied more from lower to higher compared to the one measured before loading (Figure 5). Neither the order in which the buzzes were applied and measured, nor flower age, had a significant effect on distal anther velocity or measured force in proximate anthers (Table 2).

**Table 2.** Statistical analysis of the effect of anther fusion treatment (free vs. joined), buzz order (first or second), and flower age on the RMS amplitude velocity measured in the distal anther, and RMS amplitude force experienced by the proximate anther in three *Solanum* species. Model estimates were obtained from a linear mixed-effects model with individual and/or flower identity as random effects (see Methods). Statistical significance (*P-*values) of the fixed effects was calculated with a Type III analysis of variance. Significant effects (*P-*value < 0.05) are shown in bold. The reference level used for coefficient estimation is *S. pyracanthos*, free stamen configuration.

| Velocity (mm/s; distal anther) |  |  |  |  | Force (mN; proximate anther) |  |  |
| --- | --- | --- | --- | --- | --- | --- | --- |
| Variable | Model coefficient | Estimate | SE | *P*-value | Estimate | SE | *P*-value |
|  | Intercept | 63.456 | 12.122 |  | 117.397 | 1.522 |  |
| **Treatment** |  |  |  | **<0.001** |  |  | **<0.001** |
|  | Joined | 63.273 | 8.582 |  | 6.925 | 0.795 |  |
| **Species** |  |  |  | 0.073 |  |  | **<0.001** |
|  | *S. elaeagnifolium* | -48.609 | 11.553 |  | -1.014 | 1.371 |  |
|  | *S. sisymbriifolium* | -40.225 | 10.297 |  | -7.234 | 1.263 |  |
| **Species × Treatment** |  |  |  | **<0.001** |  |  | **<0.001** |
|  | *S. elaeagnifolium* × joined | 46.774 | 12.406 |  | -4.490 | 1.149 |  |
|  | *S. sisymbriifolium* × joined | 53.146 | 10.969 |  | -4.384 | 1.016 |  |
| **Buzz order** | Order | 2.862 | 4.630 | 0.537 | -0.188 | 0.431 | 0.663 |

| Flower age | Days | -3.739 | 5.077 | 0.463 | 0.581 | 0.823 | 0.481 |
| --- | --- | --- | --- | --- | --- | --- | --- |

**Figure 4.**
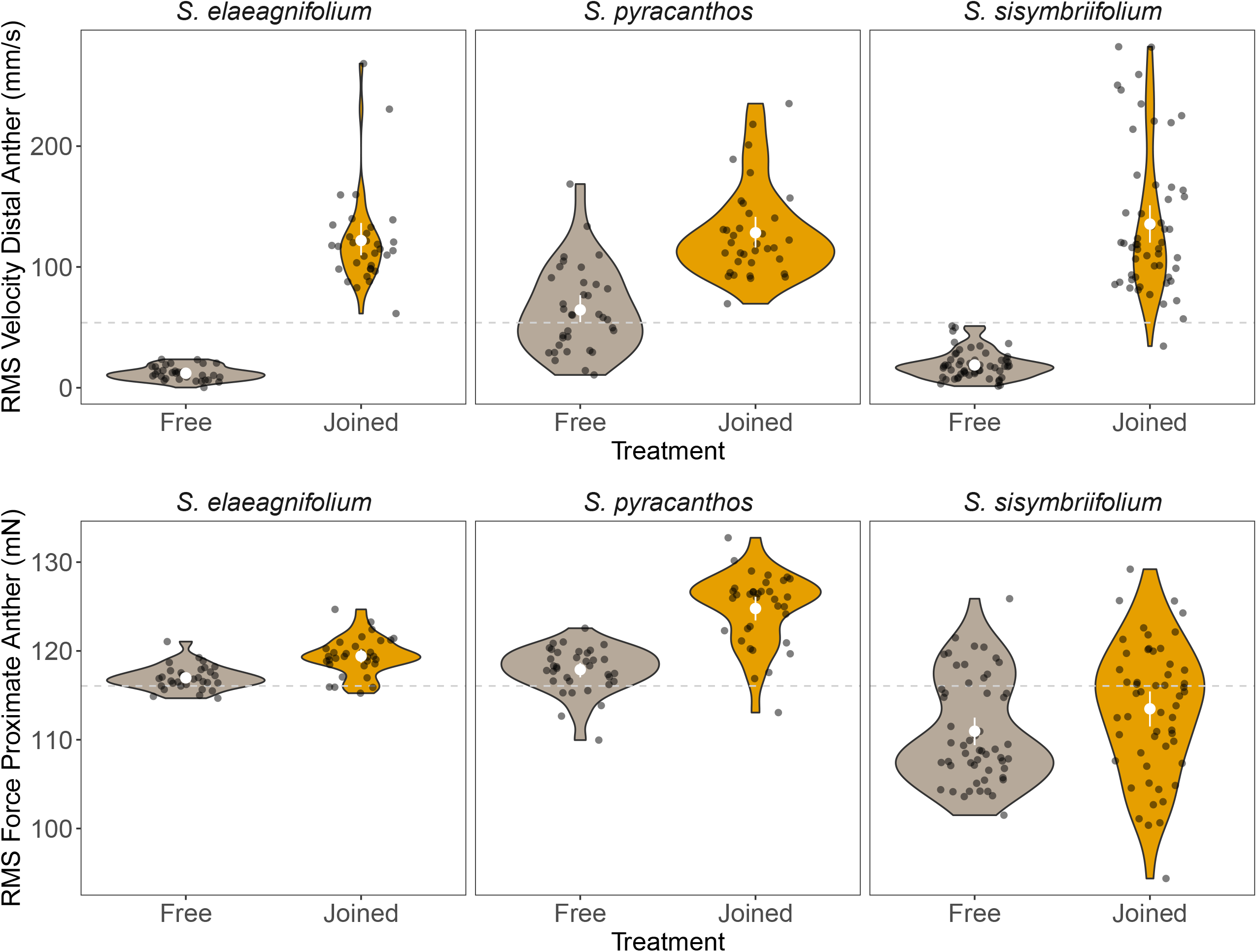
Vibration properties measured in anthers of three species of *Solanum* with experimentally joined anther cones (“Joined”; golden symbols) and without (“Free”; grey symbols). The “Joined” anther cones are created by gluing anthers together with water-based polyvinyl acetate glue (PVA). The bottom row shows the vibration force (root mean square, RMS force in mN) measured in the anther to which the mechanical vibrations were applied (proximate anther). The top row shows the vibration velocity (RMS velocity in mm/s) in a different anther in the opposite side of the cone (distal anther). As a reference, the experiment-wide median RMS values of the vibration velocity (53.85 mm/s) and force (116.05 mN) of the mechanical shaker measured both before the flower was attached to the system, and after it was removed, are shown with a dashed line. Dark circles represent observed data, with random noise added in the x-axis to facilitate visualisation. The white symbols inside the violin plots indicate the mean and 95% confidence interval of the mean calculated using 1,000 bootstrap replicates. Asterisks indicate statistical significance of within-species comparisons between treatments (marginal means contrasts): *** = P < 0.001; ** = P < 0.005.

**Figure 5.**
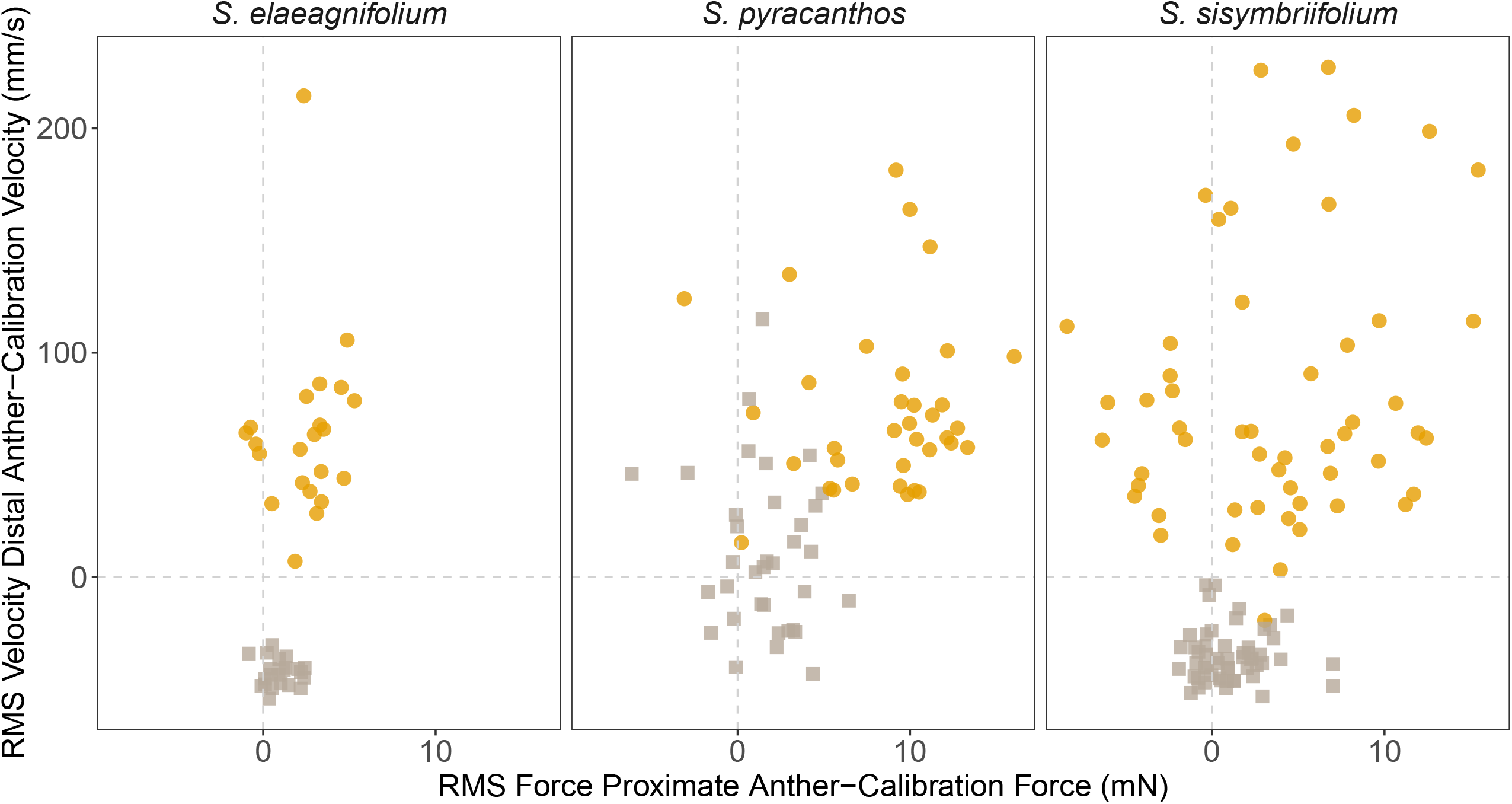
Relationship between RMS force measured in the anther to which mechanical vibrations are applied (proximate anther, x-axis) and RMS velocity of an anther located in the opposite side of the anther cone (distal anther, y-axis). RMS values are shown relative to the average daily values of RMS force and velocity measured during calibration of the mechanical shaker system. Symbol shape and colour indicate the experimental treatment: “Joined” is shown in golden circles, and “Free” in grey squares. Within each category, each symbol indicates a different flower. All flower received both treatments in random order.

Floral vibrations removed on average 9,531 ± 679 pollen grains per buzz (mean ± se; range 55-59,665, n = 240). We found a significant effect of anther treatment on pollen release that depended on plant species (species *x* treatment interaction, Table 3). For *S. elaeagnifolium* and *S. sisymbriifolium*, more pollen (1,500 – 2,300 more pollen grains; square-root transformed marginal means estimates = 48.2 ± 9.38 pollen^1/2^ and 38.3 ± 7.18 pollen^1/2^, respectively; *P* < 0.0001) was released in the joined than in the free anther treatment (Figure 6). Conversely, the average number of pollen grains released in *S. pyracanthos* was not statistically different between free and joined treatments (estimated marginal means contrast 12.3 ± 8.91 pollen^1/2^, *P* = 0.169; Figure 6). The order in which the buzzes were applied and measured did not affect pollen release, but older flowers released significantly more pollen grains (Table 3).

**Table 3.** Statistical analysis of the effect of anther fusion treatment (free vs. joined), buzz order (first or second), and flower age on number of pollen grains released after single buzzes in flowers from three *Solanum* species. Model estimates were obtained from a linear mixed-effects model with accession, individual and flower identity as random effects using a square-root transformation of pollen grains removed. Statistical significance (*P-* values) of the fixed effects was calculated with a Type III analysis of variance. Significant effects (*P-*value < 0.05) are shown in bold. The reference level used for coefficient estimation is *S. pyracanthos*, free stamen configuration.

| Variable | Model coefficient | Estimate | SE | *P*-value |
| --- | --- | --- | --- | --- |
|  | Intercept | 76.152 | 20.960 |  |
| **Treatment** |  |  |  | **<0.001** |
|  | Joined | -12.332 | 8.915 |  |
| **Species** |  |  |  | 0.609 |
|  | *S. elaeagnifolium* | -35.432 | 23.895 |  |
|  | *S. sisymbriifolium* | -47.562 | 22.725 |  |
| **Species × Treatment** |  |  |  | **<0.001** |
|  | *S. elaeagnifolium* × joined | 60.545 | 12.966 |  |
|  | *S. sisymbriifolium* × joined | 50.645 | 11.422 |  |
| **Buzz order** | Order | -4.269 | 4.842 | 0.379 |
| **Flower age** | Days | 18.777 | 6.165 | **0.003** |

**Figure 6.**
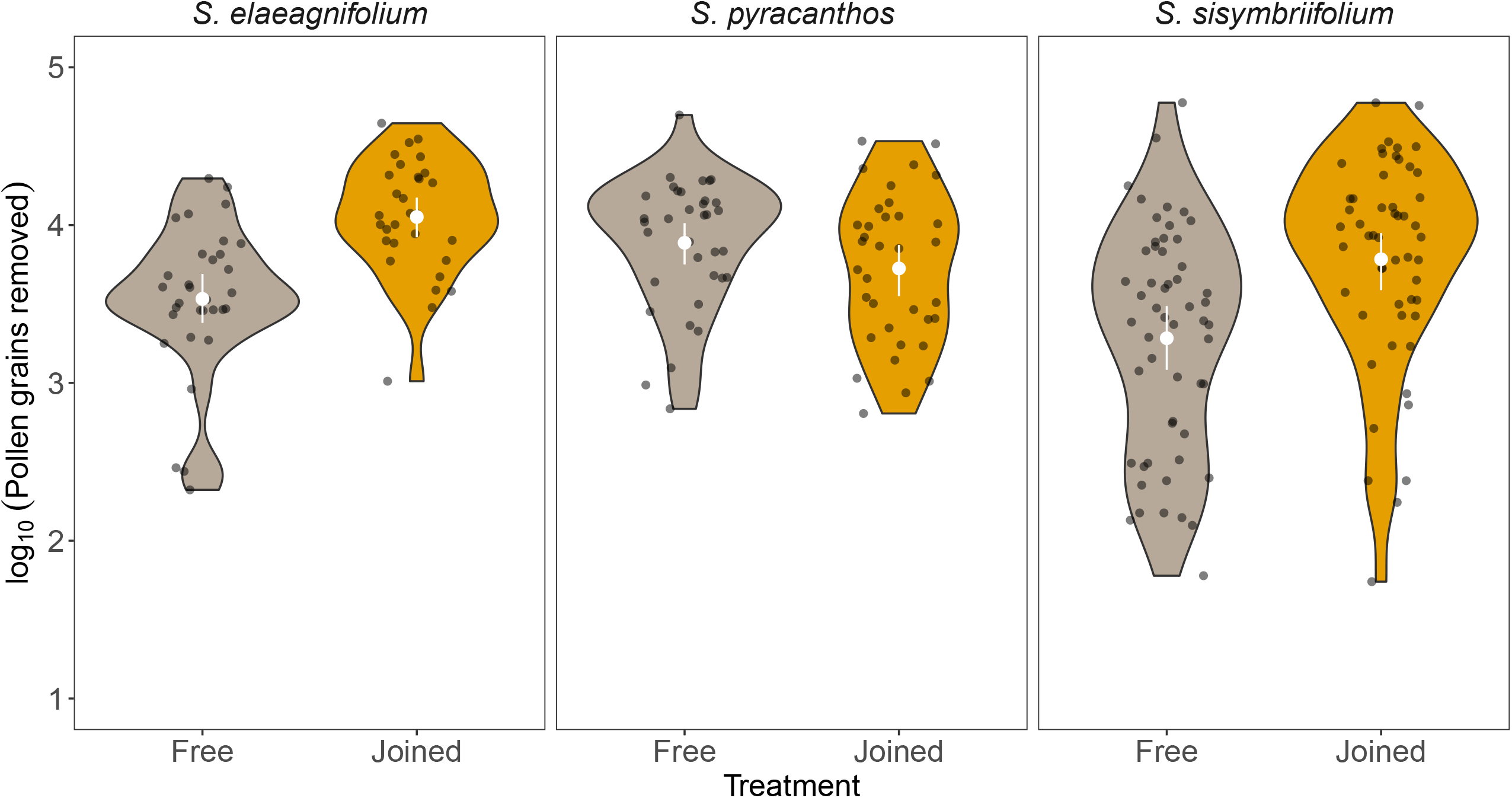
Effect of experimental manipulation of the anther cone on the number of pollen grains removed during a single buzz in three species of buzz-pollinated *Solanum* species. Dark circles represent observed data, with random noise added in the x-axis to facilitate visualisation. The white symbols inside the violin plots indicate the mean and 95% confidence interval of the mean calculated using 1,000 bootstrap replicates. Comparison of the effect of treatment (marginal means contrast): NS = P > 0.05, *** = P < 0.001.

## Discussion

In this study, we set out to address how different anther architectures, specifically the presence or absence of joined anther cones in buzz-pollinated flowers, affect patterns of vibration transmission and pollen release. Our results show that when applying vibrations to a focal (proximate) anther, the vibration velocity experienced in other (distal) anthers is significantly higher in flowers with joined anther cones, compared to those with free anthers. This result is consistent with our hypothesis that joined anther cones enable more effective transmission of applied vibrations from proximate to distal anthers via both the filament and the anther-anther pathways. Importantly, this difference in vibration transmission across anthers translates to functional differences in pollen release. In two species that naturally have loosely held, and sometimes sprawling anther architectures (*S. elaeagnifolium* and *S. sisymbriifolium*), experimental anther fusion results in more pollen released per buzz. In another species that naturally has non-joined, but tightly held anthers forming a cone (*S. pyracanthos*), experimental anther fusion did not increase pollen release. Anther fusion thus increases pollen release more strongly in species with loosely held, sprawling anthers, than in those in which anthers form a non-joined but tightly held cone. Our study suggests that anther architecture and the evolution of joined anther cones might serve to influence the rate of pollen released across the flower following vibrations applied to individual anthers.

### Functional consequences of joined anther cones

The evolution of joined anther cones is likely to incur in different costs and benefits for flower function depending on its interaction with floral visitors. Given that regardless of anther architecture, both focal and non-focal anthers release pollen upon vibrations (e.g., Nevard et al. 2021, and the present study), joined anther cones might reduce pollen wastage by directing more pollen onto the bee’s body (Glover et al. 2004). We call this the **reduced pollen wastage hypothesis**. Reducing the amount of pollen grains that misses the body of floral visitors should increase pollination success. At the same time, this anther architecture may involve a trade-off if bees can also benefit from joined anther cones by capturing a greater proportion of pollen that is released upon vibrating. For example, the stereotypical C-shape posture that bees take during the production of floral vibrations (De Luca and Vallejo-Marin 2013), should favour the receipt of pollen grains in the ventral region, where bees are generally capable of grooming and harvesting pollen grains (Vallejo-Marin et al. 2009; Huang et al. 2015; Koch et al. 2017; Tong et al. 2019). Reduced pollen wastage could also be achieved in flowers with loosely held anthers if the visiting bee can gather together all stamens using its legs and mandibles (M. Mayberry, D. McCart, J. Burrow, T.L. Ashman, and A. Russell, *unpublished data*). Bees that are relatively large compared to the flower they visit should be capable of such manipulation, although quantitative evidence of this behaviour remains scarce. At the same time, the evolution of loosely held anthers might be favored when buzzing bees can remove pollen from only one or a few anthers but only infrequently contact the stigma, such as when the bees are relatively small compared to the flower (Li et al. 2015; Solis-Montero and Vallejo-Marin 2017; Telles et al. 2020; Mesquita-Neto et al. 2021). In this context, loosely held anthers that reduce vibrations being transmitted to non-focal anthers (as shown in our study) would simultaneously reducing pollen wastage and pollen theft. We call this the **reduced costs of pollen thieves hypothesis**. The hypotheses of reduced pollen wastage and reduced costs of pollen thieves highlight how the potential benefits of fused anther cones depend on ecological context; namely the presence and abundance of floral visitors differing in size and behaviour. For example, in ecological communities dominated by large bee pollinators capable of embracing all loosely held anthers and intercepting most of the ejected pollen with their bodies (e.g. carpenter bees), the benefits of reducing pollen wastage to the plant (when pollen grains miss the visitor’s body) may be relatively minor. Conversely, the conditions for the reduced cost of pollen thieves hypothesis may prevail in communities where smaller bees acting as pollen thieves are dominant (Mesquita-Neto et al. 2018).

Additionally, the evolution of joined anther cones may increase the precision of pollen placement on specific parts of the pollinator’s body, which could increase the likelihood of stigma contact (**unimodal pollen deposition hypothesis**). During buzz-pollination, anther cones are expected to interact with the bee’s ventral side depositing pollen in a single region (unimodal), making the location of pollen placement and pick-up relatively predictable. At the same time, unimodal pollen deposition could involve a trade-off, if the pollen were thus more readily groomed into the bee’s pollen baskets (Russell et al. 2021). Accordingly, the evolution of loosely held anthers, could be favoured if pollen were thereby more frequently deposited int hard-to-groom “safe sites” on the bee’s body. The presence of loosely held anthers may also facilitate the evolution of anther specialization within a flower, if different sets of loose anthers consistently deposit pollen on different parts of the bee’s body. This preliminary division of labour may facilitate the evolution of anther morphology that improves the effectiveness of the division of labour, such as heteranthery, which commonly occurs in buzz pollinated plants (Vallejo-Marin et al. 2010; Barrett 2021). Division of labour in heterantherous species is achieved by two or more types of morphologically distinct stamens that release and deposit pollen on distinct parts of the pollinator’s body, one of which is more effectively groomed by floral visitors (feeding anthers) and another that is more likely to contact the stigma of other conspecific flowers (pollinating anthers) (Luo et al. 2008; Vallejo-Marin et al. 2009; Papaj et al. 2017; Saab et al. 2021). In heterantherous flowers, pollen release and deposition are thus expected to have a bimodal (or multimodal if more than two stamen types) distribution on the pollinator’s body, and such distribution is incompatible with joined anther cones.

Finally, a non-mutually exclusive hypothesis, for which our results provide direct empirical support, is that joined anther cones increase the transmission of vibrations across the stamens and, in some case, result in higher rates of pollen released per buzz (**increased pollen release hypothesis**). Our experiments across three *Solanum* species with slightly different anther architectures, show that in all cases, joined anther cones significantly increase the vibration amplitude (RMS velocity) transmitted to non-focal anthers showing an immediate consequence of joined anther cones. However, our experiments also show that the extent to which experimental fusion of anther cones influence pollen release is contingent on species-specific characteristics. Pollen release was increased in the two species that naturally have loosely arranged anther architectures (*S. elaeagnifolium* and *S. sisymbriifolium*), and in which the experimental treatment notably increases the extent to which anthers contact each other (the anther-anther vibration transmission pathway). An increase in pollen release through anther fusion is not observed in the Malagasy species *S. pyracanthos*, in which anthers are closely held together since the beginning of anthesis and throughout the flower’s life. Although vibration amplitude in joined anthers of *S. pyracanthos* increases, we found no effect on pollen release. This might be because, despite being non-joined by either trichomes or bio-adhesives, the closely held androecium of *S. pyracanthos* is sufficient to transmit strong enough vibrations across all anthers to maximise pollen release rate. Pollen release rate in this case might be limited not by vibration amplitude, but by other factors such as the size of the apical pore from which pollen can come out during buzzing, and the amount of freely available pollen inside the anther locules, which in part is determined by flower age (Harder and Barclay 1994; Kemp and Vallejo-Marin 2021).

The four hypotheses mentioned above are not an exhaustive list or mutually exclusive, and they could act in concert, for example by simultaneously reducing pollen wastage and increasing pollen release rates. Other consequences of free anthers could include lengthening the amount of time that a visitor spends in a flower, due to having to separately manipulate anthers, which in some cases might increase pollen deposition on the stigma, including self-pollen deposition. However, a common feature of these hypotheses is their dependence on specific ecological factors, particularly the morphological and behavioural characteristics of floral visitors. Evaluating the relative value of these hypotheses in explaining the evolution of joined anther cones, therefore requires explicitly considering the interaction of buzz pollinated plants and their wild bee pollinators.

## Acknowledgements

We thank L. Nevard and the rest of the Vallejo-Marin Lab for feedback and discussions during the development of this project, and J. Weir for support in the Controlled Environment Facility. We thank S. C. H. Barrett and A. Etcheverry for sharing their seed collection of *S. sisymbriifolium*. This research was supported by a grant from The Leverhulme Trust (RG-2018-235) to MVM.

## Supplementary Material

**Supplementary Figure 1.**
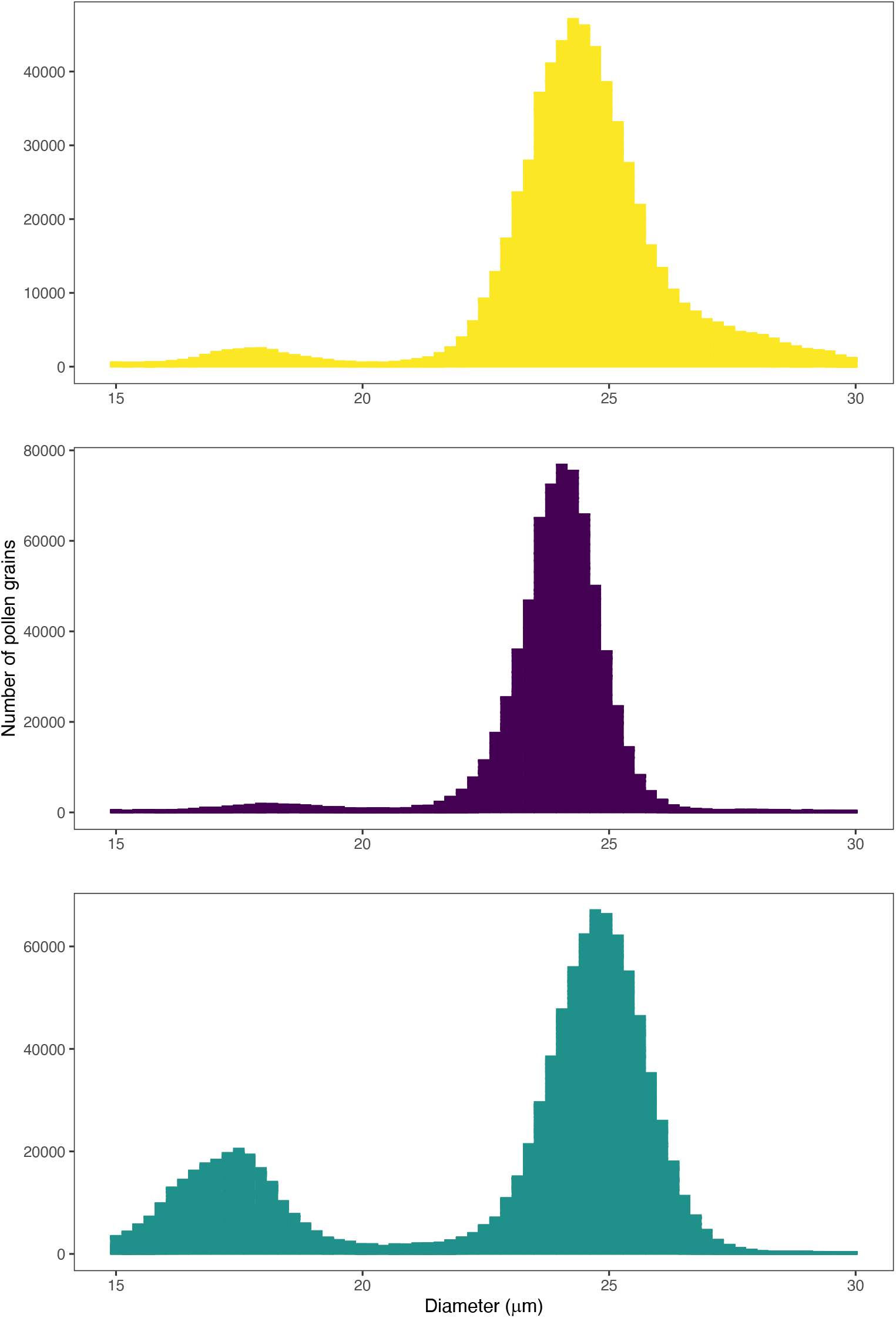
Frequency distribution of the size of measured particles (pollen grains) in the electric particle counter as described in the Methods. All samples for each species are presented in the same histogram. Top = *Solanum elaeagnifolium*. Centre = *S. pyracanthos*. Bottom = *S. sisymbriifolium*.

## Notes

### Competing Interest Statement

The authors have declared no competing interest.

### Summary of Updates

Changed the High Resolution version of Figure 3 to a Low Resolution version to reduce PDF file size.

