## Supplementary Figure 1 for "Anther cones increase pollen release in buzz-pollinated *Solanum* flowers"

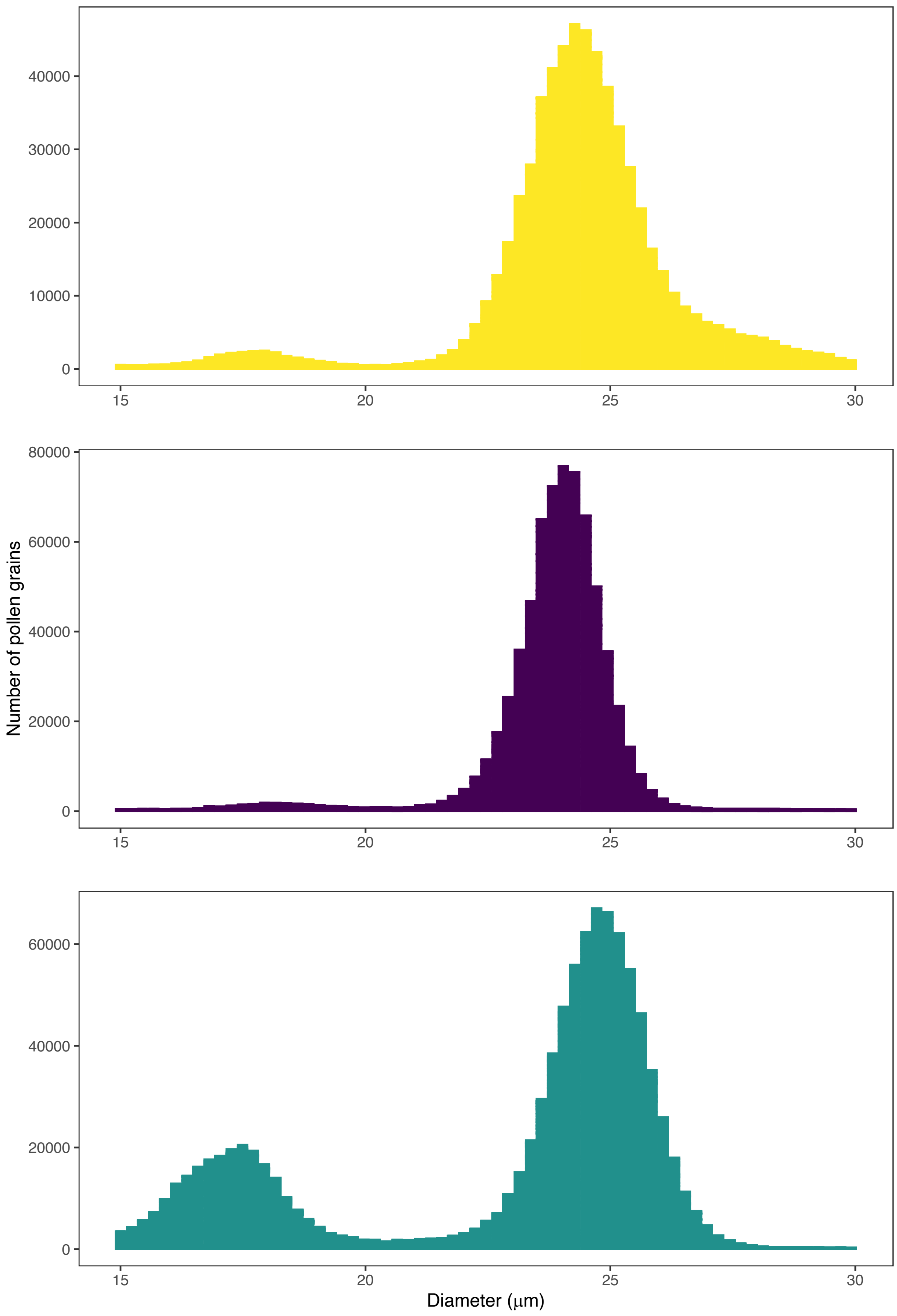
40000
30000
20000
10000
0
15
20
25
30
80000
60000
40000
20000
0
Number of pollen grains
15
20
25
30
60000
40000
20000
0
15
20
25
30
Diameter (μm)

**Supplementary Figure 1.** Frequency distribution of the size of measured particles (pollen grains) in the electric particle counter as described in the Methods. All samples for each species are presented in the same histogram. Top = *Solanum elaeagnifolium.* Centre = *S. pyracanthos.* Bottom = *S. sisymbriifolium*
